# Arabidopsis aneuploidy mutant C30-81-as6, possesses enormous variations in multiple phenotypic characteristics

**DOI:** 10.1101/2025.09.16.676461

**Authors:** Asanga Deshappriya Nagalla, Kotaro Ishii, Tomonari Hirano, Sumie Ohbu, Yuki Shirakawa, Yusuke Kazama, Tomoko Abe

## Abstract

Heavy-ion beam irradiation can be recognized as one of the powerful mutagenesis techniques because it induces high mutation frequency, minimal damage to the other traits and the possibility of having large DNA fragmental mutations. Aneuploid cells may have an additional or fewer number of chromosomes compared to their wild-type. The mutant line of C30-81-as6 is an ion beam-generated, Arabidopsis Col-0 based, aneuploid mutant line confirmed by whole genome sequence analysis, DNA quantity measurement using flow cytometry analysis and microscopic observations of additional chromosome. The mutant shows abnormal leaf shapes, petioleless rosette leaves, elevated trichome bases and later flowering phenotypes. Segregation analysis confirmed that additional DNA amount is correlated with observed phenotype since mutants showing intermediate phenotype possess moderate DNA quantities. A set of candidate causal genes was identified using up and downregulated differentially expressed genes in transcriptome analysis. Gene ontology analysis further supports mutant phenotypic characteristics and provides insights into its functional priorities, including promotion of cellular structure and energy production. Meanwhile, the downregulation of senescence-related gene ontology terms validates its extended vegetative phase. The whole genome sequence analysis based chromosome rearrangement prediction and the gene density mapping of upregulated DEGs, suggested that the long arm of chromosome 2 is a main component of the additional chromosome. In addition, the phenotype of C30-81-as6 largely resembles Arabidopsis additional chromosome 2 containing aneuploidy mutant. The ion beam mutagenesis generated aneuploidy mutant that may possess a fused DNA fragmented chromosome and it may successfully challenge the wild-type plants for establishment in nature.

## Introduction

Mutations, the changes in the DNA sequence, are essential for genetic diversity, which may ultimately lay the foundation for natural selection. Mutations can arise from various sources, including natural processes, environmental factors, and human-induced methods such as chemical mutagenesis, e.g., Ethyl Methane Sulfonate (EMS) and radiation. γ-rays, X-rays and heavy-ion beams are the common sources of radiation mutagens in Japan. The heavy-ion irradiation causes dense ionization along the particle path, resulting in various degrees of DNA damage, and is recognized as an effective mutagenic tool for biological studies and product developments including plants and microbes (Hirano et al. 2013; Ichida et al. 2008; Kanaya et al. 2008; Katano et al. 2016; Kazama et al. 2013a, 2016; Ma et al. 2013; Maeda et al. 2014; Sasaki et al. 2012; Satoh and Oono, 2019; Wang et al. 2013; Yamaguchi 2018; Yasui et al. 2012). Since the possibility of inducing a wide range of mutations, including single nucleotide variants (SNVs), deletions and chromosomal level re-arrangements (Hirano et al. 2015; Kazama et al. 2012, 2017; Naito et al. 2005). The linear energy transfer (LET) of the heavy-ions is a crucial factor in determining the type and size of the mutation induced. The LETs available for biological experiments at the RI Beam Factory (RIBF) in RIKEN range from approximately 22.5 to 4000 keV µm^-1^ (Ryuto et al., 2008). Higher-LET radiation deposits a high amount of energy that can damage the high-order structure of chromatin. It includes double-strand breaks (DSBs) that often cause errors in repairing (Hirano et al. 2013, 2021). Interestingly, the high LET can induce rare types and different sizes of mutations including large deletions, inversions, duplications and complex chromosomal rearrangements (Abe et al. 2021; Hase et al. 2017: Hirano et al. 2012, 2015; Ikoma et al. 2025; Kazama et al. 2011, 2013b, 2017; Morita et al. 2021; Oono et al. 2020; Sanjaya et al. 2021). It’s challenging to achieve these varieties of mutations with chemical mutagenesis agents such as EMS or colchicine.

Duplications and chromosomal rearrangements are types of chromosomal abnormalities that arise due to DSB followed by imprecise repair mechanisms. Ar-ion irradiation with 290 keVμm^-1^ was reported to induce a higher frequency of chromosomal-level rearrangements compared to other types of ion irradiation such as C-ions with 30 keVμm^-1^ (Hirano et al. 2012; Kazama et al. 2017). Mutant lines of Ar-365-as1 and Ar-443- as1 in the Arabidopsis background were produced by Ar-ion irradiation, confirmed to have 57.1 kbp duplicated region in Chr. 4 and 2.26 Mbp, 318.1 kbp duplicated region in Chr 2, respectively (Hirano et al. 2015). Alone with duplications, Ar-57-al1, Ar-365-as1, and Ar-443-as1 mutants revealed a total of 22 DNA fragments that have contributed to complex chromosomal rearrangements. In addition to the Ar-ions, C-ions can also induce chromosomal rearrangements, even though its frequency is relatively low, where 10.2 for Ar- ions and 2.3 for C-ions rearrangements per mutant genome, respectively. The C-ion irradiated mutant lines C30-1-as1, C30-95-as1, C30-108-as3 reported to possess 5,4 and 3 rearrangement events, respectively (Kazama et al. 2017). Furthermore, multiple rearrangements occurred in localized regions of the chromosomes, indicating the heavy-ion beam can induce clustered DNA damage (Hirano et al. 2015). In addition to Arabidopsis, a duplication event spanning about 2.5 kbp was reported in the rice Ne_50 mutant line induced by Ne ion irradiation with 31 keV μm^-1^ (Zheng et al. 2021).

DNA duplications and complex chromosome rearrangements may lead to serious chromosome abnormalities, including the existence of additional chromosomes, known as aneuploidy. The most common form of aneuploidy in humans is Down syndrome, which is caused by trisomy of Chr 21 in patients, where it’s diploid in nature. Down syndrome patients show a set of unique types of characteristics, yet they depend on the genomic and environmental variabilities (Antonarakis et al. 2004). Even though there are several genes on Chr 21 confirmed to be causal for Down syndrome, the complexity of gene dosage effects occurred not only due to the expression of genes on Chr 21, but also global gene expression patterns across the genome (Letourneau et al. 2014). On the other hand, plants have proven to be more tolerant of aneuploidy mutations. All five possible trisomics of Arabidopsis mutants are viable and have unique phenotypes (Rédei et al. 1992) and a highly heterogeneous aneuploidy population consisting of 25 different karyotypes has been previously reported (Henry et al. 2010). Those aneuploidy mutants exhibit a few key phenotypic characteristic differences compared to their euploid counterparts, yet the degree of severity of each phenotype is variable (Henry et al. 2010; Ramsey and Schemske 1998). However, an aneuploid Arabidopsis mutant derived from heavy-ion beam irradiation may not be a classical one (mutants derived from chromosome doubling agents such as Colchicine) because the responsible chromosome would be a product of duplications, complex chromosomal rearrangements and various degrees of other mutations.

In the current study, we identified a C-ion with 30 keVμm^-1^ irradiated Arabidopsis mutant that shows a series of phenotypic changes compared to wild type (WT). The pattern of inheritance of the mutant characters does not follow the Mendelian genetics ratios and flow cytometry analysis revealed that the mutant genome is significantly larger than WT genome.

The mutant cells were observed using a high-resolution microscope and an additional chromosome was observed. Due to the presence of an extra chromosome, the whole transcriptome of the mutant shows a misbehavior. The upregulated differentially expressed genes (DEGs) were densely mapped to the long arm of Chr 2. The Gene ontology (GO) analysis highlighted that the mutant is actively modifying processes related to structural components, enhancing metabolic activity while downregulating senescence-related processes. Our result provides evidence of the correlation between morphological abnormalities with DE genes and their GO terms triggered by the heterogeneous chromosomal fragments.

## Materials and Methods

### Plant materials including F_2_ population, Flowering date and phenotype observations

The *A*. *thaliana* accession Columbia-0 (Col-0) was used as WT plant. The C30-81-as6 mutant line was derived from C-ion irradiation (30 keVµm^-1^) on Col-0 dry seeds as the method described in Kazama et al. (2011). The mutant was identified in the M_2_ generation, and self-fertilized seeds from the best mutant (later flowering, crumpled, large and petiole-less rosette leaves: see the results section) were selected and grown for the next generation, up to the M_7_ generation. The F_2_ population is derived by backcrossing with Col-0, and self-fertilization of resulted F_1_ population. Seed preparation, germination, cultivation and transplanting were done based on the method described in Kazama et al. (2011). Flowering time was monitored from the days after transplanting to the soil, and growth room conditions were maintained constant throughout the experiment. The C30-81-as6 mutant plant materials used in this study were derived from the M_3_ generation unless it’s specified otherwise. To confirm the stability of the mutant trait, the best mutant was selected and grown up to the M_7_ generation.

### Microscopic observation of the chromosomes

The Chromosome observation was done using 0.4 mm diameter flower buds. The selected flower buds were fixed in the fixative solution (5:1(Methanol: 0.2M Acetic Acid)) under vacuum in 0.5 mL tube. After overnight fixation, the solution was replaced with 70% Methanol and kept overnight at 4°C. Next, the buds were washed twice with distilled water. Transfer the moisture-removed buds to an enzyme solution (Macerozyme: 0.08 g, Pectolyase: 0.02 g, Cellulase: 0.1 g and dissolve in 10 mL of acetic acid buffer (pH 4.2)) and incubate for 50 minutes in 37°C. The digested and moisture removed buds were placed on a glass slide. The buds were crushed with fine tweezers and spread using fixative solution. 7 µL of DAPI (1 mg/mL) was added to the tissues and covered with cover glass. The FV3000 Inverted Confocal Laser Scanning Microscope (Evident Corporation Tokyo, Japan) with 10 x 100 (100X/1.40 Oil UPlanSApo) magnification under 405 nm laser was used for chromosome observation.

### Flow cytometry analysis

The nuclear ploidy level was observed by using a flow cytometer (CyFlow counter, Sysmex, Kobe, Japan). Leaf samples of the same age Col-0 and C30-81-as6 mutant were collected and chopped with Otto I buffer (Otto 1990). Then the residue was filtered through a 30 µm nylon mesh filter to remove debris. The DNA staining solution (Mishiba et al. 2000) 800 µL was added to the filtrate and stained for 1 min.

### Transcriptome and GO enrichment analysis

Leaf samples of three individual plants of C30-81-as6 mutant and WT plants were collected. All the plants were the same age and C30-81-as6 mutants were confirmed as positive mutants by phenotypic observation and the presence of additional DNA content by flow cytometric analysis. The total RNA was extracted using the RNeasy Plant Mini Kit (QIAGEN, Venlo, Netherlands). The cDNA library preparation was conducted by using VAHTS Universal V8 RNA-seq Library Prep kit (Nanjing, PRC). An Illumina HiSeq/ Illumina Novaseq/ MGI2000 instrument was used for sequencing and a 2x150 paired-end configuration was adopted according to manufacturer’s instructions. Quality control was conducted using Cutadapt (V1.9.1, phred cutoff: 20, error rate: 0.1, adapter overlap: 1bp, min. length: 75, proportion of N: 0.1) (Martin 2011) to obtain high quality data. Clean reads of each biological sample were aligned to the Arabidopsis genome assembly (TAIR10) by using Hisat2 (v2.2.1) (Kim et al. 2015). The HTSeq (v0.6.1) software package (Anders et al. 2015) was used to count the reads mapped to genomic features, and to estimate gene and isoform expression levels from the pair-end clean data. Fragments Per Kilobase of transcript per Million mapped reads (FPKM) plot box distribution indicated that median values remain relatively similar among all the Col-0 and mutant samples indicating that the central tendency of gene expression levels is comparable (Fig. S1). This suggested that the occurrence of the additional chromosome hasn’t created a global expression shift and DEG analysis can be performed without additional normalizations. The resulting count matrix was used for DEG analysis using DESeq2 (Love et al. 2014) Bioconductor package, a model based on the negative binomial distribution. The DE genes were identified by adjusted *p*- value < 0.05 and a minimum two-fold change in expression. The GO enrichment analysis was conducted considering DE upregulated and downregulated genes separately using the R package clusterProfiler (v4.0) (Yu et al. 2012) with Arabidopsis thaliana annotations from org.At.tair.db (Huber et al. 2015). The Gene Ontology resource (Consortium 2021) was used as the reference database. Terms with FDR (q-value) < 0.05 are considered significant. The DEGs were mapped to the chromosomal positions using biomaRt software package (Durinck et al. 2005, 2009). The Poisson test was adapted to determine the significance of the observed number of genes in a genomic bin, followed by Benjamini-Hochberg (BH) procedure to control the False Discovery Rate (FDR).

### Whole-Genome Mutation Analysis and Linkage Analysis

A single positive M_3_ mutant and C30-81-as6-77 mutant plants were selected and the DNA was extracted from leaf samples using DNAeasy plant minikit (QIAGEN, Venlo, Netherlands), with using IDTE 1×TE solution pH 8.0 (Integrated DNA technologies, Iowa, USA). The DNA libraries were created using MGIEasy PCR-Free DNA library prep kit (MGI Tech, Shenzhen, China). The whole genome sequencing was performed using MGI DNBSEQ-G400 instrument (MGI Tech) in paired-end, 2 × 150-bp mode. Bioinformatics analysis was conducted using AMAP (Ishii et al. 2016) implemented on the HOKUSAI-BigWaterfall supercomputing system (RIKEN). The AMAP includes programs for quality control, mapping, detection of various degrees of mutation and creating a final analysis integrating the analysis results. The resulting candidate mutations were visually confirmed using IGV software (Robinson et al. 2011).

## Results

### The C30-81-as6 with delayed flowering exhibits a range of phenotypic variations in leaves, roots, and trichomes

The M3 plants of C30-81-as6, identified as a mutant line with abnormal leaf morphology, exhibited multiple phenotypic alterations, including delayed flowering, which occurred approximately 5 weeks after transplantation into soil, in contrast to the wild-type plants that flowered within about 1 week (Table 1). The leaf-related abnormal characteristics included crumpled, large and petiole-less rosette leaves (Fig. 1). The mature leaves are large, wide, thick and fleshy with clear sinuate edges compared to WT mature leaves. This mutant is delayed in flowering and has an increased number of rosette leaves during the long vegetative period (Table 1 and Fig. 1). We grew inbred populations up to the M_7_ generation to check the stability of mutant characters. The M_7_ mutants have resembled phenotypic features, including a delay of flowering time (Table 1).

**Fig. 1.**
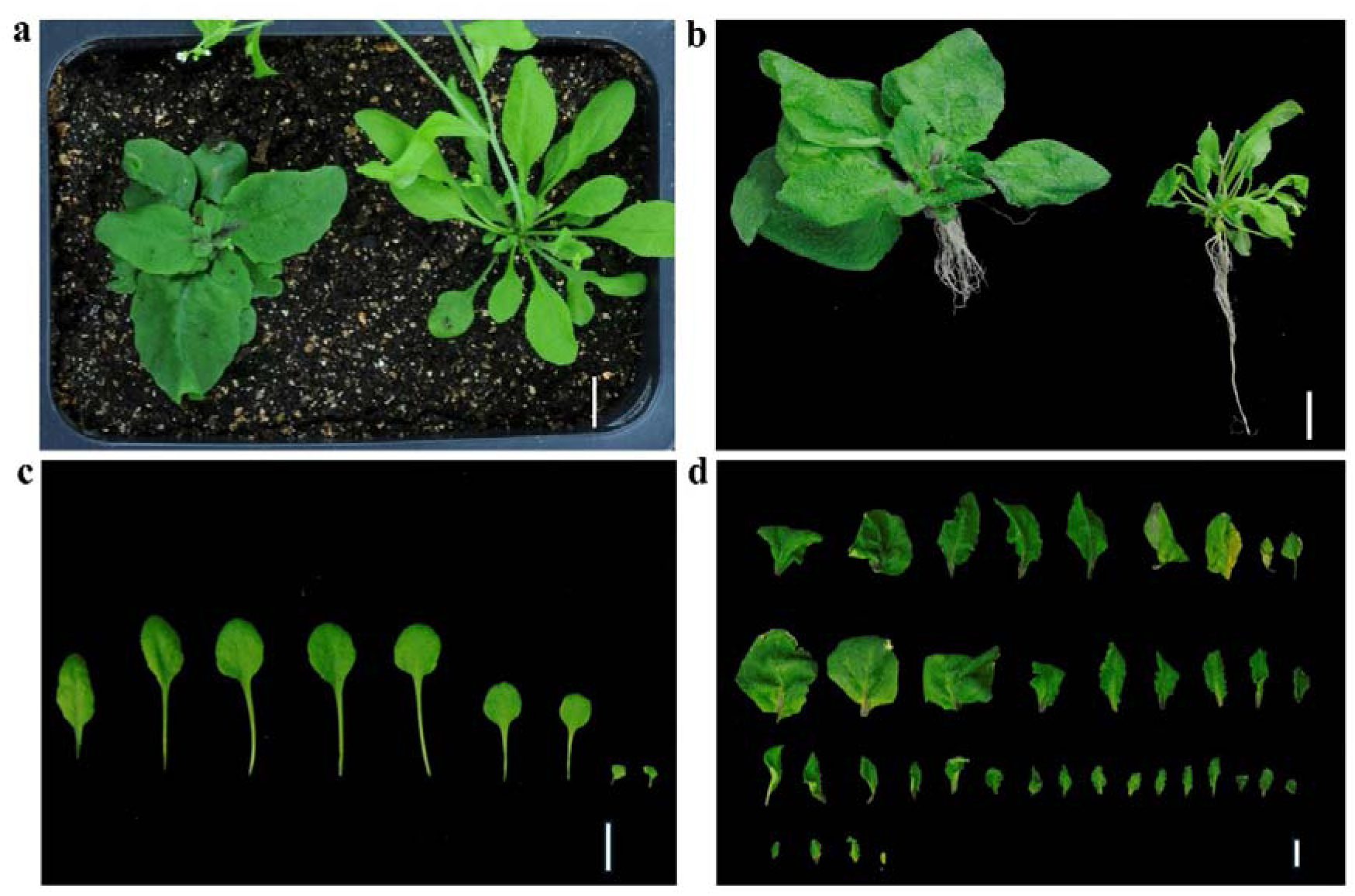
Phenotypic characteristics of the C30-81-as6 mutant and WT plants. (a) The mutant (left) flowered late, with large and petiole-less rosette leaves compared to the WT (right). (b) The mutant possesses a shallow root system (left) compared to the WT (right). The leaf panel of WT (c) and mutant (d) shows the differences in leaf morphology and the number of leaves. Photos were taken when the plant showed flowering WT: around 7 days after transplant and C30-81-as6: around 35 days after transplant. Bar = 1cm.

**Table 1.** Flowering time of the C30-81-as6 mutant and WT plants. Data are means ± SD (n ≥ 4).

| Line | M<sub>3</sub> | M<sub>7</sub> |
| --- | --- | --- |
| Col-0 | 7.1 ± 0.9 | 8.6 ± 1.0 |
| C30-81-as6 | 35 ± 2.8 * | 32.3 ± 5.6 * |

\* The significance of the difference was assessed by Student's t-test (\* $P$ < 0.01).

The leaves of the C30-81-as6 mutant appeared to have a rougher surface than those of the WT. Initially, we suspected that this was due to an increase in trichome density. However, microscopic observations revealed that there was no significant increase in trichome density; instead, the trichome bases were elevated. The trichome bases of the mutant exhibited a mountain-like shape. In contrast, those of the WT appeared flat (Fig. 2). In addition to leaf morphology, the C30-81-as6 mutant showed interesting root structural abnormalities compared to the WT. The mutant had a shallow and dense rhizosphere, whereas the WT had a relatively deep and thin rhizosphere (Fig. 1).

**Fig. 2.**
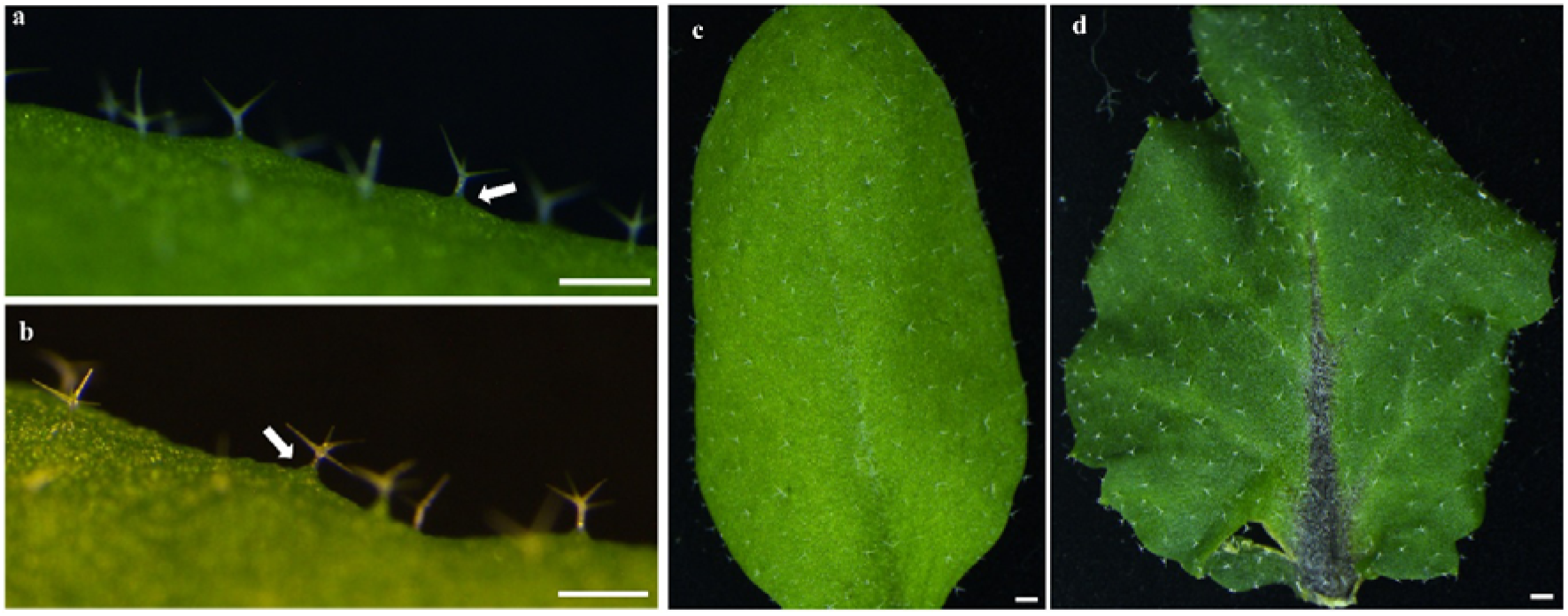
Trichome base and leaf base morphology of the C30-81-as6 mutant and WT plants. The trichome base of the Col-0 (a) is relatively flat but the mutant (b) has a heap-shaped trichome base. The white color arrows point to the trichome base. The leaf surface of the Col- 0 (c) and the mutant (d). Bar = 0.5 cm.

### The C30-81-as6 mutant does not follow the Mendelian genetic ratios of inheritance

There were various types of segregants in the M_3_ population. Approximately 3.3% exhibited all of the above phenotypic characteristics and were designated as “positive mutants”. If a mutant does not satisfy at least a single character: later flowering, crumpled, large and petioleless rosette leaves were considered as negative mutants. To obtain a homozygous population, we grew several inbred populations. The best positive mutant was selected and its seeds to grow the next population. The number of positive mutants was 9.0%, 25.0% and 36.8% in the M_4_, M_5_ and M_7_ generations, respectively (Table S1). Therefore, the occurrence of the positive mutants was low in consecutive inbred populations, indicating the possibility of multiple genes responsible for the positive mutant phenotype.

### Identification of the gene responsible for the phenotypic change in the C30-81-as6 mutant

To investigate the responsible gene/s for the phenotypic variations in the C30-81-as6 mutant, first, we tried WGS analysis. The automated mutation analysis pipeline (AMAP; Ishii et al. 2016) resulted in six SNVs, one deletion, and 13 chromosomal breakpoints, including intra-chromosomal translocations (ITX), and inter-chromosomal translocations (CTX). Totally, nine homozygous mutated genes were found (Table S2). The mutations were confirmed by manual observation in Integrative Genomics Viewer (IGV; Robinson et al. 2011). We performed a linkage analysis using an F_2_ population of 84 individuals and only one plant (C30-81-as6-77) represented an intermediate phenotype of all three mutant characteristics: later flowering, crumpled large leaves and petioleless rosette leaves. The number of positive mutants was (the single intermediate plant) insufficient to perform linkage analysis. Therefore, we consider individuals showing at least two mutant characters. There were 22 plants showing later flowering, 21 plants showing crumpled, large leaves and one plant showing petioleless rosette leaves. Altogether, 13 plants showed both later flowering with crumpled large leaves and only one plant showed all three mutant characteristics (Fig. S2). We collected DNA from those 14 plants and used it for linkage analysis. This linkage analysis resulted none of the nine targets being genetically linked with at least a single mutant phenotype.

### Increased DNA content and identification of additional candidate chromosomes in the C30-81-as6 mutant

We performed flow cytometry analysis to measure the genome size of the mutant using leaf samples of Col-0 and C30-81-as6 plants. The histogram indicates four clear peaks of nuclear DNA content. The 2C peak representing diploid nuclei shows a slight shift in DNA content in the mutant. Importantly, the 4C, 8C and 16C peaks represent the tetraploid, octoploid and hexaploidy nuclei showing a clear shifting of DNA content in the mutant compared to Col-0, suggesting significantly increased genomic DNA levels (Fig. 3). The ratio of DNA content of Col-0 and C30-81-as6 for each peak is 1.1 (2C:1.09, 4C:1.12, 8C:1.12, 16C:1.12). This data on the additional DNA content reflected the possibility of having an additional chromosome in the mutant genome.

**Fig. 3.**
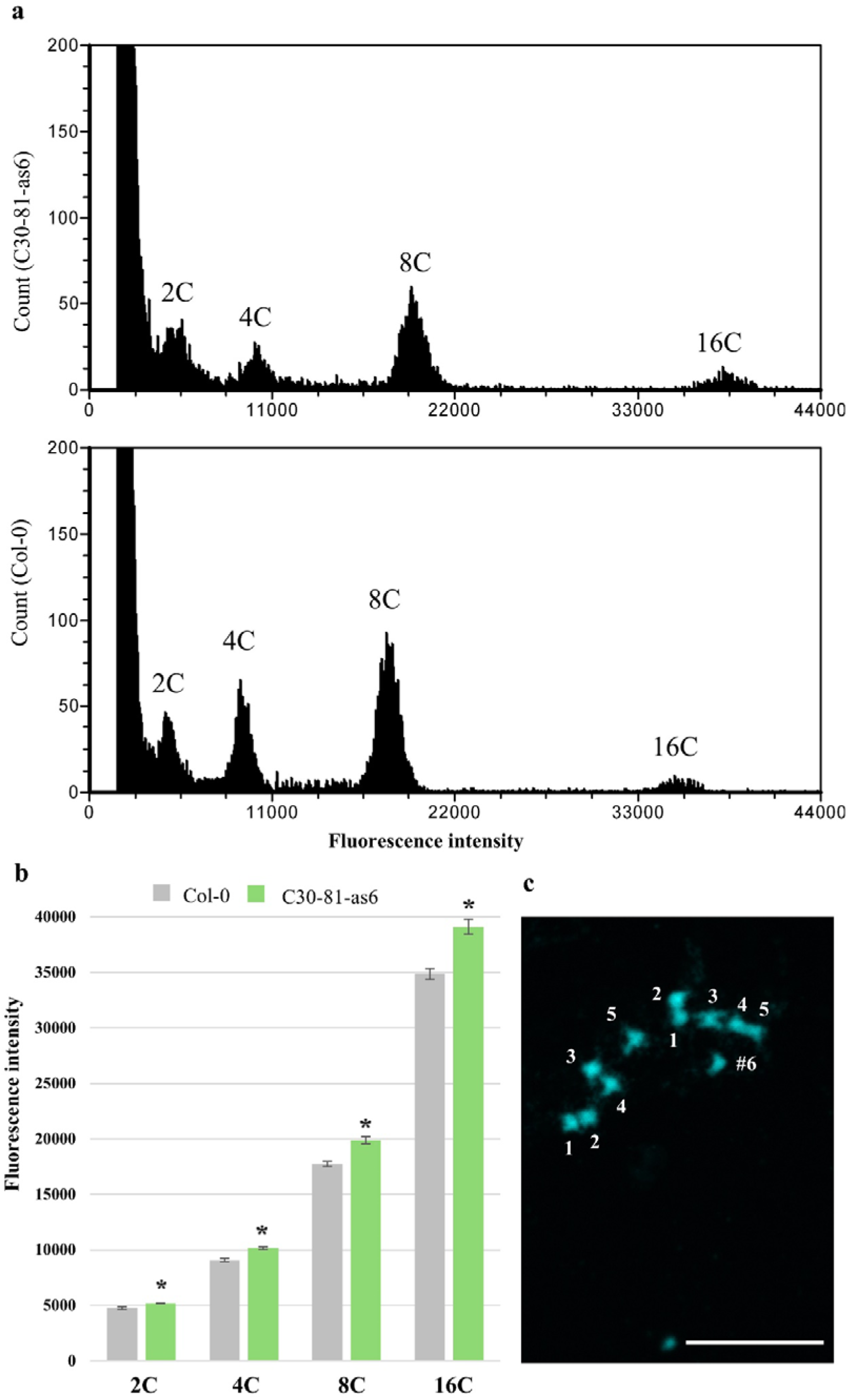
Flow cytometry analysis in C30-81-as6 mutant and Col-0 plants. (a) Histogram of fluorescence intensity of the nuclei in mutant (upper) and Col-0 (below). 2C, 4C, 8C and 16C peaks represent the diploid, tetraploid, octoploid and hexaploid nuclei, respectively. (b) Comparison of the fluorescence intensity for each peak. The significance of the difference was assessed by Student’s t-test (*P < 0.01) and n ≥ 3. (c) Microscopic observation of chromosomes. Chromosome observation of the cells from flower bud samples of C30-81-as6 mutants in M_3_ generations under 1000X magnification. The numbers 1-5 and #6 represent the number of chromosomes. Bar = 10 μm. Chromosomes observation was performed twice using flower buds from independent generations and obtained the same results.

To ascertain this possibility, we observed the cell nuclei of M_3_ and M_7_ (photo not shown) mutants (additional DNA content was verified by flow cytometry analysis for each mutant plant) flower bud samples. Under 1000× magnification, clear chromosomes were observed in the candidate cells undergoing division. There were five and six chromosomes containing cells, confirming the positive mutants are consistent with an additional chromosome named #6 (Fig. 3c).

To elucidate potential chromosome rearrangements, fragment breakpoints and rejoined junctions were visualized using WGS data analysis. A reciprocal, CTX between Chr. 1 and Chr. 2 resulted including a tandem duplication and a deletion in the second repeat in Chr. 2 (mutation ID 1 and 2 in Fig. 4a). Since the Chr. 1 side is homozygous and the Chr. 2 side is heterozygous (mutation ID 2), it is considered that there is another copy of Chr 2 that does not carry the translocation (Table S2). The same CTX was observed in the WGS analysis performed using C30-81-as6-77 genomic DNA (Fig. S3 and Table S3), indicating this translocation could be responsible for key mutant characters. Another candidate reciprocal CTX was reported between Chr. 3 and Chr. 4. However, the exact position on the Chr. 3 side is unknown due to the occurrence of the repetitive region (mutation ID 8). An intra-chromosomal translocation with an inversion was observed in Chr. 5, although the correct positions are unknown due to repetitive sequences (mutation ID 12 and 13). Furthermore, upregulated DEGs were mapped against the position of each chromosome (details about the transcriptome analysis are mentioned below). Interestingly, many upregulated DEGs were significantly mapped to the long arm of Chr 2 with higher density compared to other chromosomes (Fig. 4b). Therefore, both chromosomal rearrangement and DEG mapping analysis propose that the long arm of Chr 2 would be a major causal fragment for the mutant phenotype. Since we were unable to confirm the genetic linkage with other homozygous mutated genes with the mutant phenotype, the presence of the CTX in Chr.1- Chr. 2 and the additional chromosome would be the key mutations responsible for C30-81-as6 phenotypic characters.

**Fig. 4.**
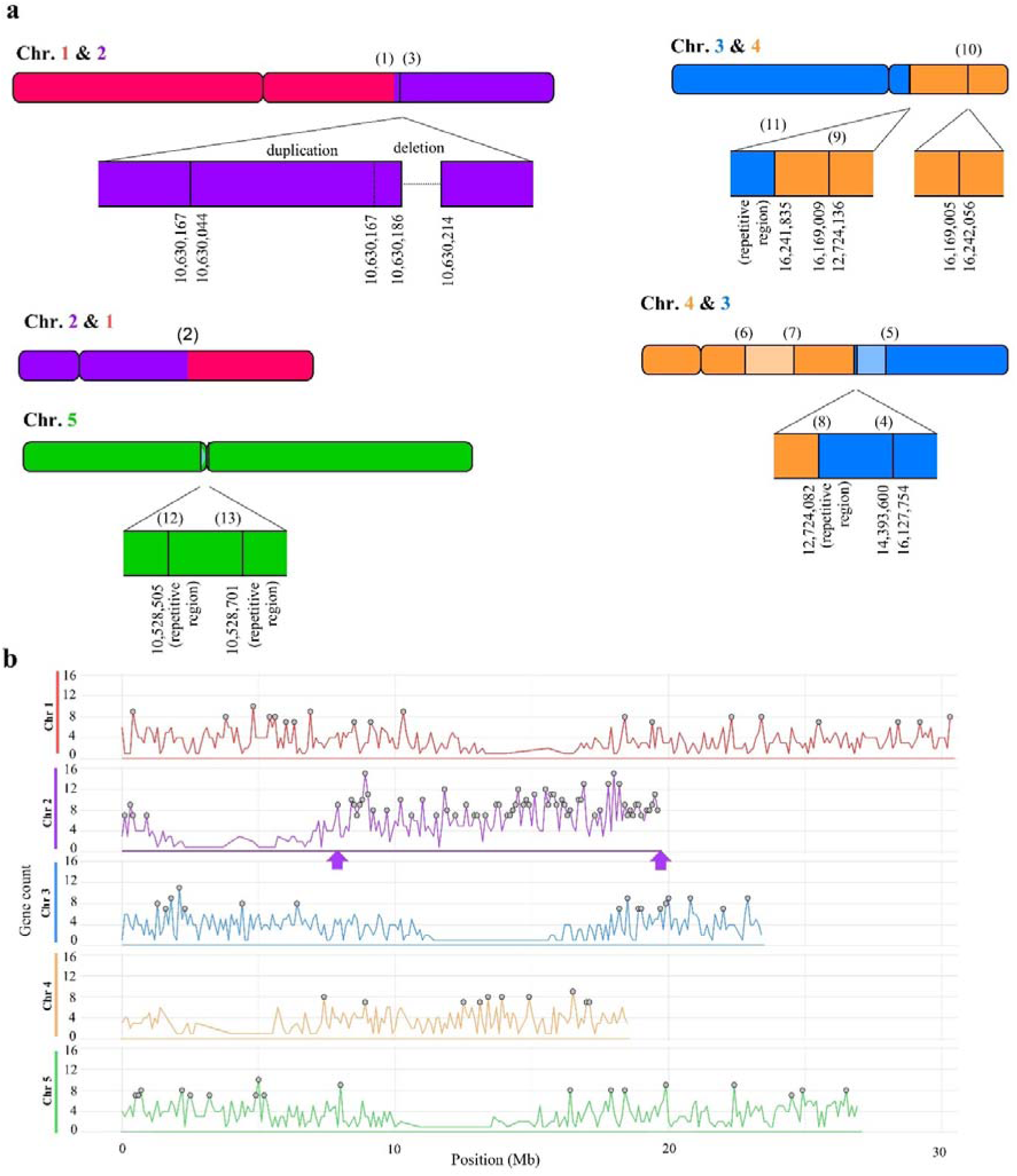
Visualization of additional chromosome fragments (a) diagram of proposed reconstructed chromosome structures of the C30-81-as6 mutant based on the Automated Mutation Analysis Pipeline. The mutation IDs (1-13) represent the candidate break\rejoin locations as indicated in Table S2, based on TAIR 10. Pale-colored fragments indicate those rejoined in an inverted direction and the sizes of the chromosome fragments are proportionate to the actual nucleotide sizes. (b) Gene density plot of positions of the upregulated DEGs on each chromosome. Bin size =100 Kb. Gene count refers to the number of genes in each bin and the red color circle indicates the statistically significant gene count based on the Poisson test (*p* adj < 0.05). The arrows indicate the candidate chromosome fragment expected to be presence of higher number of upregulated DEGs due to duplication.

### Segregation analysis of C30-81-as6 mutant

Throughout the M_3_-M_7_ generations, we found plants showing intermediate characteristics, including leaf shape and flowering time. Next, we were interested in understanding the correlation of additional DNA content in controlling mutant phenotypic characters. In a situation of partial aneuploidy where the additional chromosome is expected to be compaction and segmental duplications occur in meiosis, it’s possible to obtain plants with intermediate DNA content (DNA amount is greater than WT, but less than aneuploidy mutant: C30-81-as6) (Henry et al. 2005, Henry et al. 2010). Therefore, we checked the DNA content of the intermediate phenotype display plants, mutants and WT using flow cytometry. As expected, the early flowering plants show weak leaf phenotypes (shape and size) and less additional DNA compared to positive mutants. When comparing the DNA content of several mutants that flower between the earliest-flowering wild type and the latest-flowering mutant (positive mutant), the increase in DNA content was greater in the later-flowering mutants (Table 2 and Fig.5). These results suggest that the additional DNA quantity is associated with the overall mutant phenotype.

**Fig. 5.**
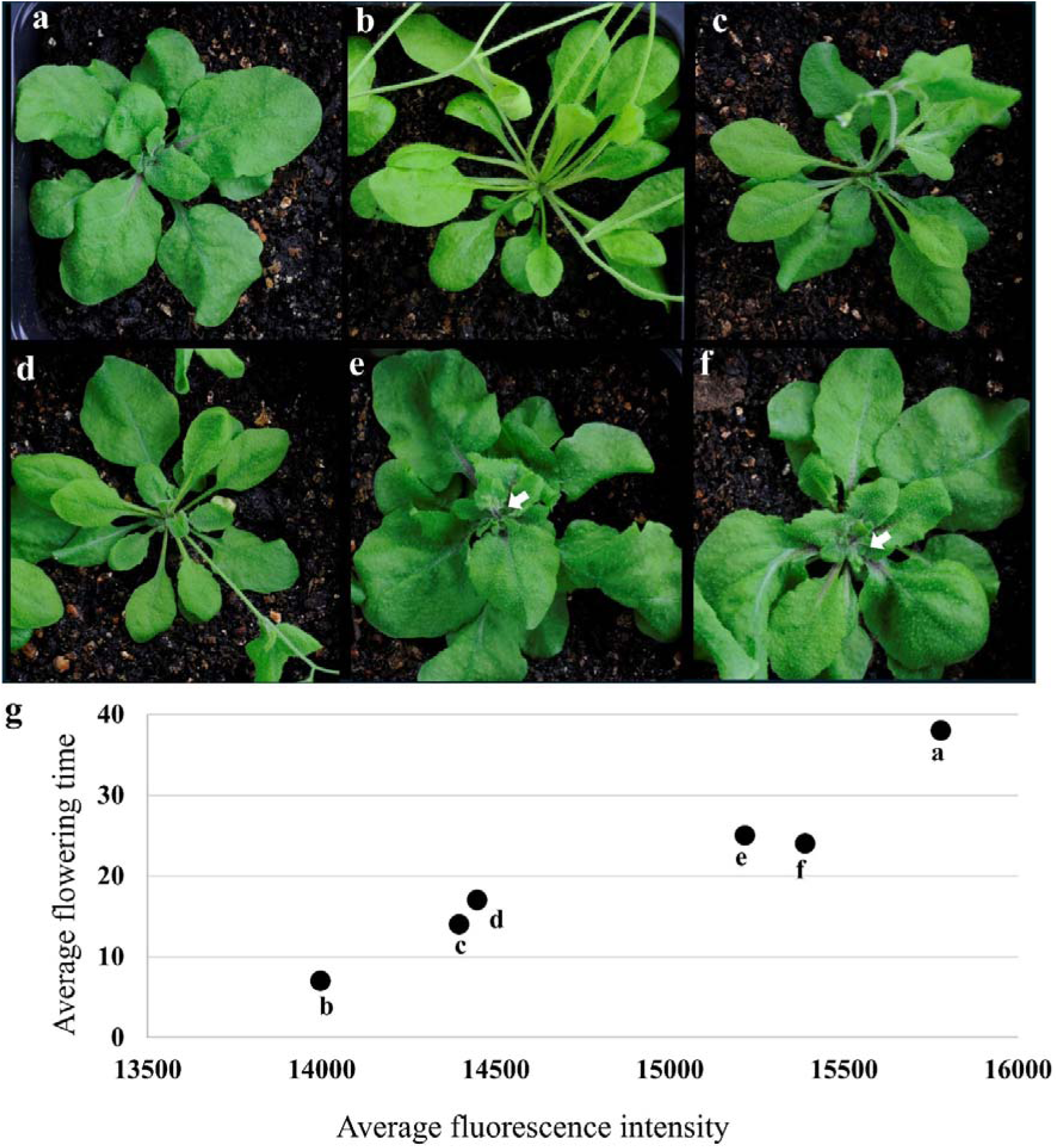
The plants with intermediate additional DNA content showing weaker phenotype characters compared to C30-81-as6. (a) C30-81-as6: positive mutant. (b) WT. (c,d) C30-81- as6 mutants showing weak leaf shape and very early flowering compared to positive mutants. (e,f) C30-81-as6 mutants showing strong leaf shape and later flowing compared to WT. (g) Flowering time delaying with increment of fluorescence intensity in each mutant (a-f). White arrows indicate the emergence of immature flower buds (around 25 days after transplant).

**Table 2.**
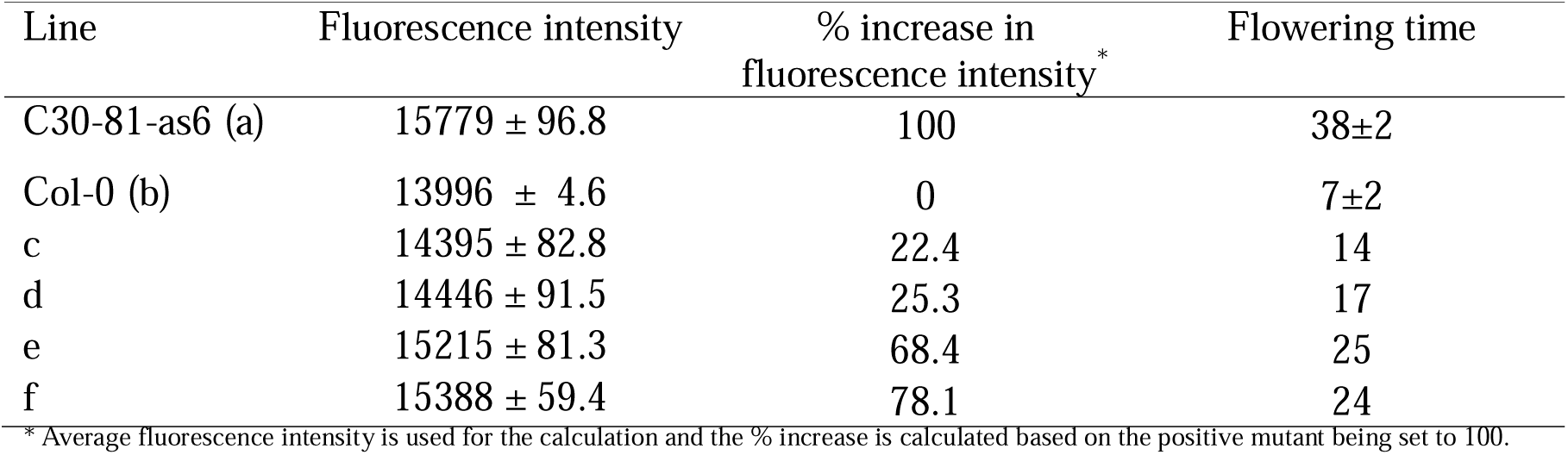
Fluorescence intensities of the selected peak and their % increment represent the additional DNA quantities of C30-81-as6 plants. Symbols (a-f) resemble symbols in Fig .5. Data are means ± SD (n ≥ 3). For (a, b) data from individual plant leaves (biological replicates) and (c-f) data from three leaves from the single plant (technical replicates) were measured.

| Line | Fluorescence intensity | % increase in fluorescence intensity * | Flowering time |
| --- | --- | --- | --- |
| C30-81-as6 (a) | 15779 ± 96.8 | 100 | 38±2 |
| Col-0 (b) | 13996 ± 4.6 | 0 | 7±2 |
| c | 14395 ± 82.8 | 22.4 | 14 |
| d | 14446 ± 91.5 | 25.3 | 17 |
| e | 15215 ± 81.3 | 68.4 | 25 |
| f | 15388 ± 59.4 | 78.1 | 24 |

\* Average fluorescence intensity is used for the calculation and the % increase is calculated based on the positive mutant being set to 100.

### The C30-81-as6 mutant possesses an extremely misregulated transcriptome compared with WT

Based on the above results, the mutant shows a series of phenotypic deviations due to the segregation of an additional chromosome. Because of the appearance of phenotypic defects, we hypothesized that the mutant’s transcriptome would be severely misregulated mainly due to the over-expression of a cluster of genes positioned on the duplicated regions . To understand the impact on the whole transcriptome, we performed a transcriptome analysis using the positive mutants confirmed by flow cytometry analysis and Col-0 plants. A total of 4096 upregulated and 3022 downregulated DEGs were identified (Fig. 6). The total number of DEGs accounted for about 30% of the whole transcriptome. The GO analysis for upregulated genes emphasized that the most significant and the highest number of genes are involved in plant-type cell wall biogenesis, mitotic cell cycle and ribosome biogenesis and as the key biological processes (BP), structural molecular activity, including structural constituents of ribosome and cytoskeleton protein binding as the key molecular functions (MF) and Cellular Component (CC) of the most genes are ribosome, ribosomal subunit, and cytosolic ribosome. The GO terms for downregulated genes resulted that, the highest number of genes with significance recorded in BP are plant organ senescence, leaf senescence and response to decreased oxygen levels. Salt transmembrane transporter activity, metal ion transmembrane transporter activity and UDP-Glycosyltransferase activity are results under the top three GO terms for MF. The downregulated DEGs which functionally connected to CC are plasma membrane-bounded cell projection, cell projection and pollen tubes (resulted only three GO terms). However, the number of genes identified in CC is quite low compared to BP and MF (Fig. 6). Collectively, the GO terms for upregulated genes outline the mutants’ prioritization for structural components production, energy and precursor molecule acquisition. Interestingly, the plant senescence regulatory genes are strictly downregulated in the mutant, providing the genetic basis for the extended vegetative phase supported with structural components produced by upregulated genes.

**Fig. 6.**
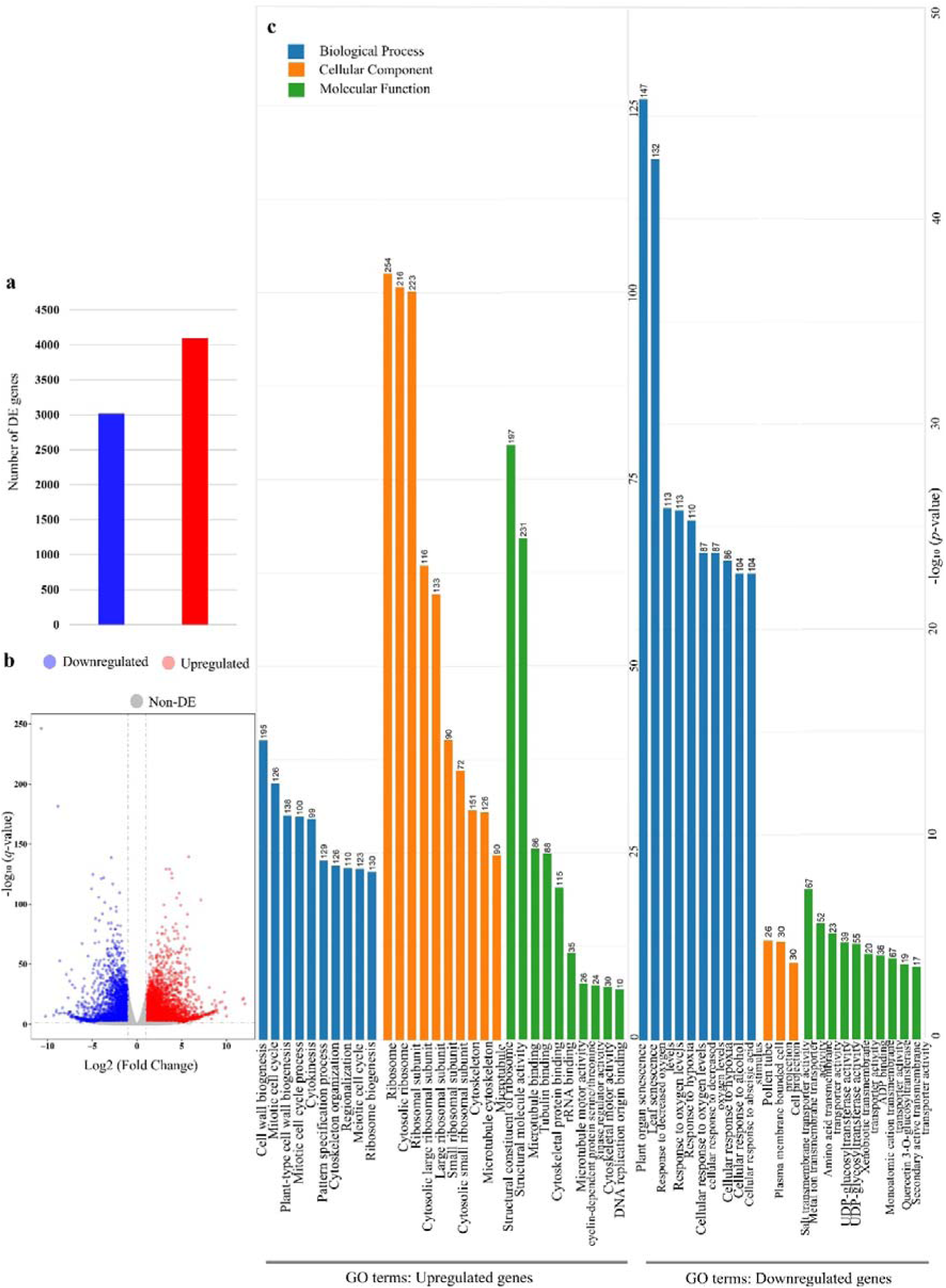
Overview of transcriptome analysis results of C30-81-as6 mutants and Col-0. (a) the number of up and downregulated DEGs in the mutant compared to Col-0. (b) volcano plot showing the distribution of significantly up and downregulated DEGs. (c) a summary of gene ontology (GO) terms for up and downregulated DE genes. The top 10 statistically significant GO terms show in the bar plot and the number in front of the bar represents the number of genes supported for the corresponding GO term.

## Discussion

### Increased DNA content, additional chromosome and the mutant genome

Based on the flow cytometry results and the microscopic observation of the chromosomes, we were able to confirm that the C30-81-as6 plants possess an additional chromosome. In the mutant, half of the cells are predicted to have five chromosomes, and the other half are expected to have six chromosomes, resulting in a DNA content ratio of C30-81-as6/WT of exactly 1.1 (Table 2 and Figure 3b). However, we did not find any mutants with DNA ratios exceeding this value. When this mutant produces gametes by meiosis followed by self-fertilization, there can be five possible chromosome arrangements in the zygote. **1.** 5 (n)+5 (n) – **10 (2n)**: WT type zygote. **2.** 5 (n)+6 (n) – **10 (2n)+1 (n)**: aneuploid heterozygous type zygote. **3.** 6 (n)+6 (n) – **12 (2n)**: aneuploid homozygous type zygote. **4.** 5 (n)+7 (n) – **10 (2n)+ 2 (2n)**: aneuploid zygote (if one chromosome is lost due to misegregation or if an extra chromosome is retained due to nondisjunction). **5.** 6 (n)+7 (n) – **10 (2n)+3 (n)**: aneuploid zygote (due to misegregation or nondisjunction). The DNA ratio of 1.1 is assumed to match the 2^nd^ type of plant above. If the plant is produced from the 3^rd^, 4^th^, or 5^th^ types of zygote, the DNA ratio should be ≥1.2. Since we could not find such a plant, the 3^rd^, 4^th^, or 5^th^ type of zygote was assumed to be an embryonic lethal. The additional chromosome fragments are anticipated to be produced through the fusion of duplicated and rearranged chromosome fragments. Therefore, in a zygote of 12 (2n), there might be four copies of certain genes, and those genes would be extremely upregulated, leading to lethal conditions. Even though the actual C30-81-as6 mutant is a type of heterozygous plant, it is the only existing mutant with phenotypic characters. Furthermore, we identified segmental aneuploid mutants, which exhibited intermediate traits regarding flowering time (Fig. 5) and DNA content, falling between the wild type and the positive mutants, suggesting a correlation between increased DNA content and delayed flowering in these mutants (Fig. 5).

The number of positive mutants was continuous, yet at a slow rate, increasing in the consecutive inbred populations (Table S1). One of the possible reasons for this situation would be chromosomal instability leading to more 10 (2n) +1 (n) offspring. If 10 (2n)+1 (n) plants consistently produce n = 6 gametes more frequently than expected, then crosses with n = 5 gametes will continuously regenerate 10 (2n)+1 (n) offspring at an increasing rate. Another possible reason can be identified as the biased chromosome transmission where one type of gamete (n = 6 gametes) is preferentially produced over the n = 5 gametes.

### Transcriptome analysis reveals the candidate genes responsible for the mutant phenotype

An additional chromosome is expected to be responsible for increasing the number of copies of a gene and possibly the gene may be overexpressed. However, based on the transcriptome data, both upregulated and downregulated DEGs resulted (Fig 6). This may be due to the feedback transcriptional regulation, indicating that the overexpressed and downregulate genes that are responsible for the opposite biological function. Transcriptome data provide clues to explain the mutant abnormal phenotypic features. The C30-81-as6 is a flowering time-related mutant and both downregulation of important floral promoters and upregulation of floral repressors were reported. The floral promoters include *FLOWERING LOCUS T* (*FT*), *GIGANTEA*, *Flavin-Binding -Kelch Repeat-F-Box 1* (*FKF1*) and *FRUITFULL* (*AGL8*), which were downregulated in the mutant. Meanwhile, upregulation of floral repressors *TERMINAL FLOWER 1* (*TFL1*), *CURLY LEAF* (*CLF*), *VERNALIZATION 5* (*VRN5*) and *AGAMOUS-LIKE 15* (*AGL15*) were observed in the mutant. Simultaneously, C30-81-as6 has abnormalities related to leaf shape, leaf development and petiole development. The differentially expressed *CURLY LEAF* (*CLF*), *Growth-regulating factor 2* (*AtGRF2*), *AUXIN RESPONSE FACTOR 7* (*ARF7*), *BLADE ON PETIOLE2* (*BOP2*) and *KANADI 1*(*KAN1*) are promising candidate genes which possibly responsible for those abnormal phenotypes (Table 3).

**Table 3.**
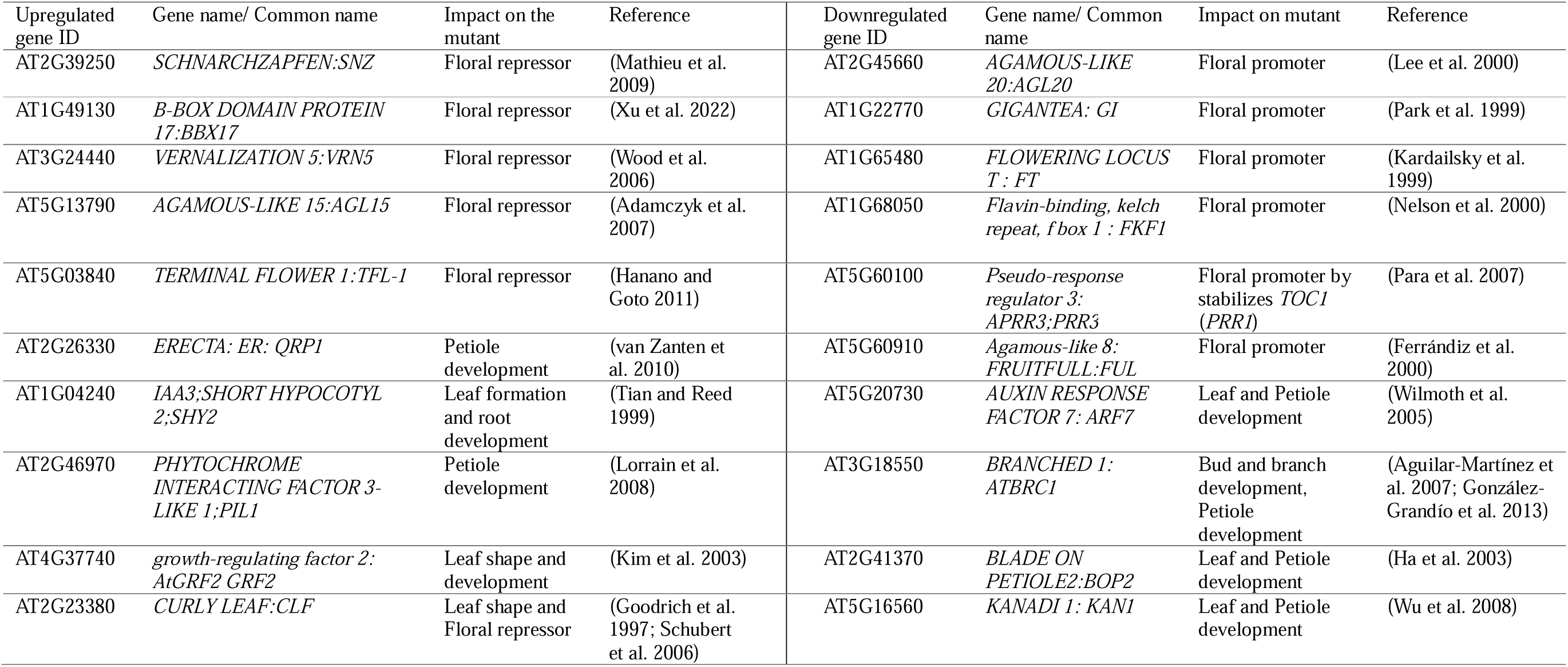
Important DEGs and their candidate impact on the C30-81-as6 mutant.

The GO terms represent an interesting overview of the whole transcriptome behavior of the mutant. Based on the BPs of the upregulated genes, the mutant cells are actively remodeling their cell wall structure, increasing cell division and growth through the mitotic cell cycle while ramping up protein production (via ribosomes). These biological processors create a relationship with GO terms of MF because the upregulated genes are involved in the functions of structural integrity, protein synthesis machinery, and cytoskeletal organization. The GO terms for CC further support the theme of enhanced protein synthesis capacity in the mutant. Because the highest number of upregulated genes are located related to ribosomes. Collectively, the GO terms of upregulated genes correlate the mutant phenotype characteristics of extended vegetative phase, larger leaves, and increased number of leaves. Because the mutant cells are assembling ribosomes, using them to synthesize structural proteins, and directing those proteins to active cell division, remodeling the cell wall and cytoskeleton. The additional chromosome in the mutant likely disrupts regulators that balance vegetative growth and flowering. Downregulation of plant organ senescence, leaf senescence, and response to decreased oxygen levels (hypoxia response) in the BP category of GO analysis is consistent with the C30-81-as6 phenotype. Because the mutant prioritizes vegetative growth by suppressing processes that limit growth or promote aging. Downregulation of the GO terms of MF related to salt and metal ion transmembrane transporter activity suggests that the mutant sacrifices stress resilience mechanisms for prioritization of growth. Because the downregulation of transmembrane related genes may reduce the capacity to manage salt stress or ion balance (Wu et al. 1996). Only three GO terms with low gene counts related to CC resulted in the analysis. Some of the downregulated DE genes may be functionally associated with cellular structures like root hairs and trichomes. Interestingly C30-81-as6 mutant possesses phenotypic abnormalities in the root and trichome structures (Fig. 1 and Fig. 2). Root hairs are plasma membrane-bounded projections extending from epidermal cells and trichomes are also cell projections that rely on cytoskeletal regulatory genes. Previous reports suggest that root hair mutants show defects in genes regulating cell projection growth (Galway et al. 1999; Yuen et al. 2005). Therefore, abnormal shallow root formation and trichome bases suggest defective cell projection formation, possibly due to downregulated CC terms. In summary, highly statistically significant up and downregulated GO terms are likely correlate with mutant phenotypic features. As a whole, the mutant extends its vegetative phase by delaying flowering and increasing the biomass. The GO terms may expose the mechanisms that provide clues such as prioritizing the production of cell mass, structural components, and protein production while suppressing organ senescence.

### Arabidopsis Chromosome 2 may be a major component of the additional chromosome in the C30-81-as6 mutant

Classically, colchicine treatment has been used to obtain polyploid cells in Arabidopsis. It disrupts the microtubule formation and longer exposures can inhibit entire cytokinesis (Henry et al. 2005; Parra-Nunez et al. 2020; Parra[Nunez et al. 2024). In addition, previous results have proven that chromosomes appear in triplicate, are vulnerable to break and rearrange (Huettel et al. 2008). In contrast, C30-81-as6 is a product of heavy-ion beam irradiation, which is a physical mutagenesis method. The mutant possesses a single additional chromosome, but two additional chromosome-containing plants were absent in the population. However, we have not yet identified the exact composition of the additional chromosome. The WGS mutation analysis-based chromosome rearrangement suggested a fragmental translocation between Chr. 1 and Chr. 2. The paired ends of Chr.1 and Chr.2 resulted as a homozygous and heterozygous, respectively (mutation ID 1 and 2 in Fig 4a and Table S2). In addition to this reciprocal translocation, a heterozygous ITX was found in Chr. 2, suggesting the possibility of having a native copy of the long arm of chromosome 2. However, heterozygous ITX in mutation ID 3 can occur even with just two copies of the chromosome (one chromosome has ITX and the other one Wt). Alternatively, it supports 3 copies of chromosomes (two chromosomes homozygous ITX and one Heterozygous ITX or the opposite way). Microscopic observation and DNA quantity measurements support the presence of 3^rd^ chromosome. Therefore, we propose that the occurrence of the 3 copies of Chr. 2 as the most appropriate model.

Upregulated DEGs were densely mapped to the long arm of Chr 2 since there can be three copies of those genes and it might be a part of the additional chromosome. The previous study Henry *et al*. 2010 compared the phenotypic consequences of inducing combinations of aneuploidy in Arabidopsis (Henry et al. 2010), and the aneuploidy mutant (Figure 1c in Henry et al. 2010) shows a resemble phenotype with C30-81-as6 mutant (Fig. 1 and Fig. 5a). Surprisingly, its karyotype is 2n + 1 chromosome 2. This data confirms the fact that the additional chromosome in the C30-81-as6 mutant may possess a large fragment of Chr 2. Furthermore, the resemble phenotype supports the concept that similar phenotypes can be observed in individuals having the same karyotype (Henry et al. 2010). However, the impact on the phenotype varies from one individual to another, possibly due to external factors such as growth conditions. In the case of C30-81-as6, since it’s an ion-beam induced mutant line, additional mutations may also modify the phenotype. Mutations are prone to occur in the natural context, as a part of the evolutionary pathway. Even though aneuploidy or chromosome-level mutations have a low probability and often disadvantageous in mutation breeding, it has the potential to induce outstanding phenotypic modifications. When some of the characteristics including leaf shape, vegetative to reproductive phase transition and root structures match beneficially, the mutant can thrive in nature. C30-81-as6 mutant represents a best-candidate phenotype model for this kind of circumstance.

## Data availability

Nucleotide sequence data files are available in the NCBI Sequenced Read Archive under the accession number SUB15363077 and SUB15611655 (https://www.ncbi.nlm.nih.gov/sra/PRJNA1271611) for C30-81-as6 and C30-81-as6-77, respectively. The transcriptome data files are available in the NCBI Gene Expression Omnibus under the accession number GSE299045 (https://www.ncbi.nlm.nih.gov/geo/).

## Supporting information

Supplemental files for Arabidopsis aneuploidy mutant C30-81-as6 MS

## Acknowledgments and Funding

We thank the RIKEN Nishina Center and the Center for Nuclear Study, the University of Tokyo for operating RIBF for performing the ion-beam irradiation. We are grateful for the technical help the Support Unit for Bio-Material Analysis, RIKEN CBS Research Resources Division provides regarding Sanger sequencing. We thank RIKEN R-COMS for lending us the FV3000 microscope and technical support. The bioinformatics analysis was performed using the HOKUSAI-BigWaterfall supercomputing system (RIKEN, Saitama, Japan) under project numbers Q22208 and Q23443. We would like to thank Dr. Ryouhei Morita for supporting lab experiments.

## Author Contributions

Study conception and design: Tomoko Abe, Yusuke Kazama, Asanga Deshappriya Nagalla. Performed experiments and data analysis: Asanga Deshappriya Nagalla, Tomonari Hirano, Sumie Ohbu, Yuki Shirakawa. Performed WGS analysis and transcriptome analysis: Kotaro Ishii, Asanga Deshappriya Nagalla. Funding acquisition: Tomoko Abe. Visualization: Asanga Deshappriya Nagalla, Tomonari Hirano. Writing: Asanga Deshappriya Nagalla. Review and editing: Asanga Deshappriya Nagalla, Yusuke Kazama, Kotaro Ishii, Tomonari Hirano, Tomoko Abe.

## Statements and Declarations

Conflict of interest: The authors declare that they have no known competing financial interests or personal relationships that could have appeared to influence the work reported in this paper.

