## Supplemental files for Arabidopsis aneuploidy mutant C30-81-as6 MS for "Arabidopsis aneuploidy mutant C30-81-as6, possesses enormous variations in multiple phenotypic characteristics"

**The affiliations of the authors:**

1. RIKEN Nishina Center, 2-1 Hirosawa, Wako, Saitama 351-0198, Japan

2. Department of Radiation Measurement and Dose Assessment, Institute for Radiological Science, Quantum Life and Medical Science Directorate, National Institutes for Quantum Science and Technology, Chiba, Japan,

3. Faculty of Agriculture, University of Miyazaki, Miyazaki, Japan,

4. Department of Bioscience and Biotechnology, Fukui Prefectural University, Eiheiji-cho, Japan

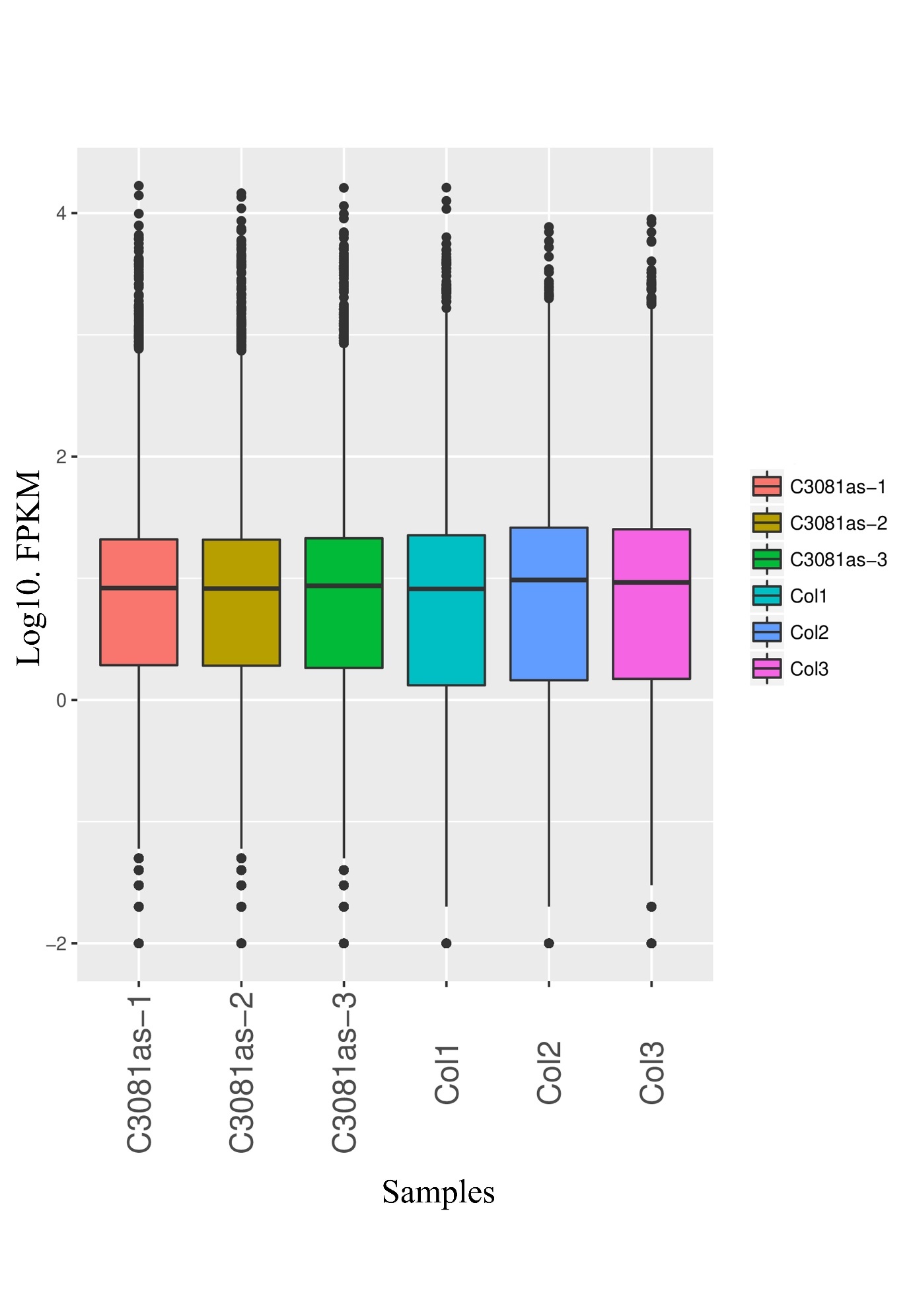

**Supplementary Fig. 1**

Fragments per kilobase of transcript per million mapped reads (FPKM) plot box distribution of RNA-seq data visualize the distribution of gene expression levels across samples.

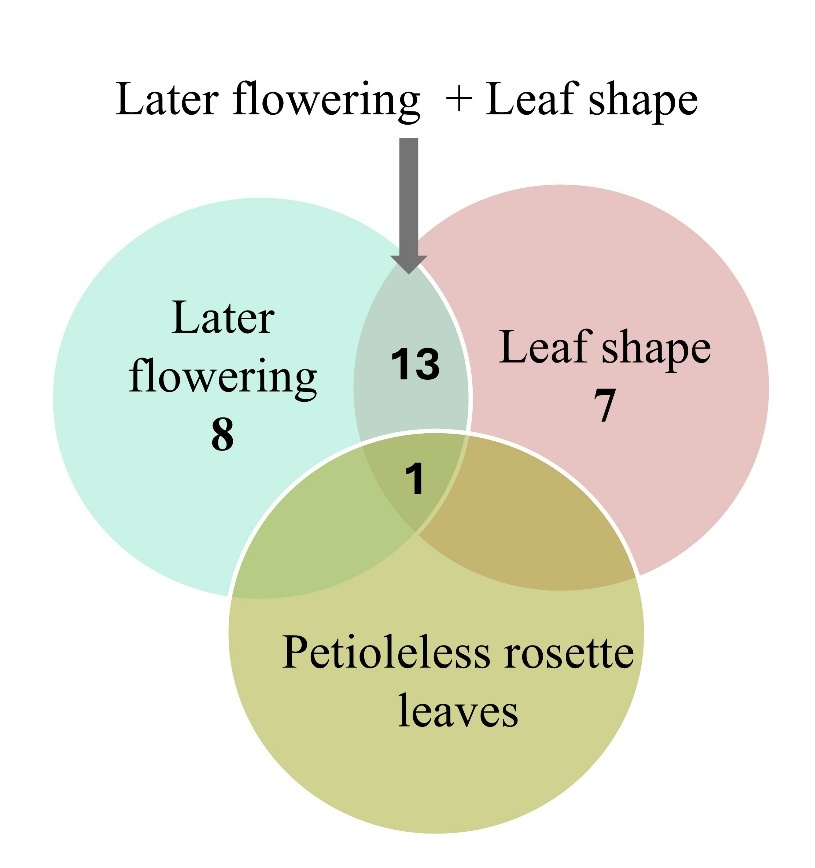

**Supplementary Fig. 2**

The phenotypic characters and the number of plants observed in F_2_ population.

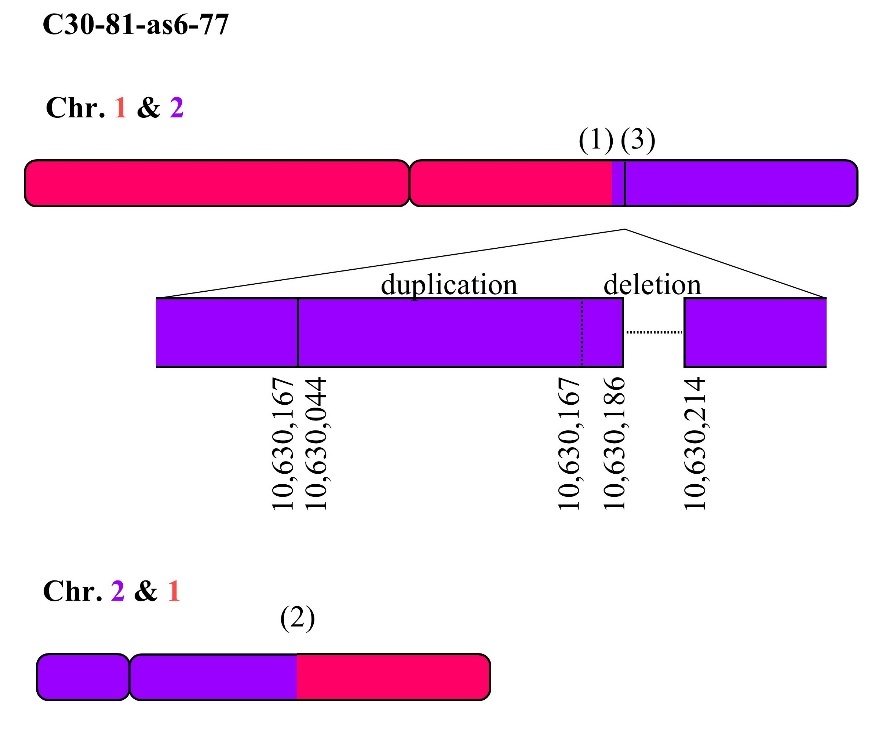

**Supplementary Fig. 3**

Diagram of proposed reconstructed chromosome structures of the C30-81-as6-77 mutant based on the Automated Mutation Analysis Pipeline. The mutation IDs (1-3) represent the candidate break\rejoin locations as indicated in Table S3, based on TAIR 10. The sizes of the chromosome fragments are proportionate to the actual nucleotide sizes.

**Supplementary Table 1**: The number of positive mutants (later flowering, crumpled, large and petioleless rosette leaves containing plants) in each generation.

| Generation | Total number of plants | Positive mutants | % of positive mutants |
| --- | --- | --- | --- |
| M_3_ | 30 | 1 | 3.3 |
| M_4_ | 33 | 3 | 9.0 |
| M_5_ | 40 | 10 | 25.0 |
| M_7_ | 19 | 7 | 36.8 |

**Supplementary Table 2:** Whole genome sequence analysis results of C30-81-as6 mutant.

| Chromosome 1 | Position 1 | Genetic homogenity | Chromosome 2 | Position 2 | Genetic homogenity | Reference | Mutated base | Type of mutation | Effect type for gene | Gene ID affected by position 1 | Gene ID affected by position 2 | Mutation ID |
| --- | --- | --- | --- | --- | --- | --- | --- | --- | --- | --- | --- | --- |
| 1 | 1,279,595 | **HOM** |  |  |  | C | A | SNP | missense_variant | AT1G04610 |  |  |
| 1 | 3,150,930 | **HOM** |  |  |  | G | A | SNP | synonymous_variant | AT1G09730 |  |  |
| 1 | 7,285,221 | HET |  |  |  | C | A | SNP |  |  |  |  |
| 1 | 10,085,618 | HOM |  |  |  | G | A | SNP |  |  |  |  |
| 1 | 11,639,575 | HOM |  |  |  | AT | A | DEL |  |  |  |  |
| 1 | 17,740,018 | HOM |  |  |  | T | C | SNP |  |  |  |  |
| 1 | 18,557,695 | HOM |  |  |  | A | G | SNP |  |  |  |  |
| 1 | 20,690,572 | HOM |  |  |  | T | A | SNP |  |  |  |  |
| 1 | 22,888,469 | **HOM** | 2 | 10,120,869 | **HET** |  |  | CTX | trancation |  | AT2G23770 | **(1)** |
| 1 | 22,888,477 | **HOM** | 2 | 10,120,491 | **HET** |  |  | CTX | trancation |  | AT2G23770 | **(2)** |
| 1 | 26,521,774 | HOM |  |  |  | AT | A | DEL |  |  |  |  |
| 1 | 28,293,670 | HOM |  |  |  | GA | G | DEL |  |  |  |  |
| 2 | 653,478 | HET |  |  |  | T | A | SNP | missense_variant | AT2G02470 |  |  |
| 2 | 653,480 | HET |  |  |  | C | A | SNP | missense_variant | AT2G02470 |  |  |
| 2 | 9,992,666 | HET |  |  |  | TTG | T | DEL |  |  |  |  |
| 2 | 10,630,044 | HET | 2 | 10,630,167 | **HET** |  |  | ITX |  |  |  | **(3)** |
| 2 | 10,630,186 | HET |  |  |  |  | 27 bp DEL | DEL |  |  |  |  |
| 2 | 10,958,868 | HET |  |  |  | A | C | SNP |  |  |  |  |
| 2 | 11,263,038 | HET |  |  |  | T | A | SNP |  |  |  |  |
| 2 | 14,154,316 | HET |  |  |  | A | T | SNP | splice_donor_variant&intron_variant | AT2G33400 |  |  |
| 2 | 16,930,529 | HOM |  |  |  | AC | A | DEL |  |  |  |  |
| 2 | 19,205,018 | HOM |  |  |  | AT | A | DEL |  |  |  |  |
| 3 | 1,709,369 | **HOM** |  |  |  | C | T | SNP | stop_gained | AT3G05760 |  |  |
| 3 | 5,528,102 | **HOM** |  |  |  | C | T | SNP | missense_variant | AT3G16310 |  |  |
| 3 | 7,581,255 | HOM |  |  |  | C | T | SNP |  |  |  |  |
| 3 | 11,719,665 | HET |  |  |  | C | A | SNP |  |  |  |  |
| 3 | 11,738,017 | HET |  |  |  | C | CA | INS |  |  |  |  |
| 3 | 14,315,102 | HET |  |  |  | C | T | SNP |  |  |  |  |
| 3 | 14,393,600 | HET | 3 | 16,127,754 | HET |  |  | ITX | trancation |  | AT3G44540 | **(4)** |
| 3 | 14,393,875 | HET | 3 | 16,127,768 | HET |  |  | ITX | trancation |  | AT3G44540 | **(5)** |
| 3 | 14,711,314 | HET |  |  |  | A | G | SNP |  |  |  |  |
| 3 | 16,235,901 | HET |  |  |  | TTC | T | DEL |  |  |  |  |
| 3 | 18,217,794 | HET |  |  |  | A | T | SNP | missense_variant | AT3G49142 |  |  |
| 3 | 19,710,233 | HET |  |  |  | G | C | SNP | splice_region_variant&intron_variant | AT3G53180 |  |  |
| 3 | 19,879,576 | HET |  |  |  | G | T | SNP |  |  |  |  |
| 3 | 22,702,225 | **HOM** |  |  |  | A | C | SNP | missense_variant | AT3G61340 |  |  |
| 3 | 23,052,522 | HOM |  |  |  | A | T | SNP |  |  |  |  |
| 3 | 23,360,253 | **HOM** |  |  |  |  | 138 bp DEL | DEL | frameshift_variant | AT3G63230 |  |  |
| 4 | 347,586 | HOM |  |  |  | T | C | SNP |  |  |  |  |
| 4 | 2,913,481 | HOM |  |  |  |  | 30 bp DEL | DEL |  |  |  |  |
| 4 | 6,153,345 | HET | 4 | 9,111,121 | HET |  |  | ITX |  |  |  | **(6)** |
| 4 | 6,153,351 | HET | 4 | 9,111,129 | HET |  |  | ITX |  |  |  | (7) |
| 4 | 8,928,591 | HET |  |  |  | G | A | SNP |  |  |  |  |
| 4 | 9,298,550 | HET |  |  |  | G | C | SNP |  |  |  |  |
| 4 | 10,709,765 | HET |  |  |  | C | A | SNP |  |  |  |  |
| 4 | 11,911,295 | HOM |  |  |  | A | G | SNP |  |  |  |  |
| 4 | 12,724,082 | **HOM** | 3 | (repetitive region) |  |  |  | CTX | trancation | AT4G24660 |  | **(8)** |
| 4 | 12,724,136 | **HOM** | 4 | 16,169,009 | **HET** |  |  | CTX | trancation | AT4G24660 |  | **(9)** |
| 4 | 16,169,005 | HET | 4 | 16,242,056 | HET |  |  | CTX |  |  |  | **(10)** |
| 4 | 16,241,835 | HET | 3 | (repetitive region) |  |  |  | CTX |  |  |  | **(11)** |
| 4 | 15,826,641 | HET |  |  |  | G | T | SNP |  |  |  |  |
| 4 | 15,826,642 | HET |  |  |  | A | T | SNP |  |  |  |  |
| 4 | 16,359,953 | HET |  |  |  | GA | G | DEL |  |  |  |  |
| 5 | 613,920 | HET |  |  |  | C | A | SNP | synonymous_variant | AT5G02720 |  |  |
| 5 | 613,921 | HET |  |  |  | G | GTTGTACACAAA | INS | frameshift_variant | AT5G02720 |  |  |
| 5 | 10,528,505 | **HOM** | 5 | (repetitive region) |  |  |  | ITX |  |  |  | **(12)** |
| 5 | 10,528,701 | **HOM** | 5 | (repetitive region) |  |  |  | ITX |  |  |  | **(13)** |
| 5 | 16,740,452 | HET |  |  |  | A | C | SNP |  |  |  |  |
| 5 | 16,867,074 | HET |  |  |  | C | CGG | INS | frameshift_variant | AT5G42210 |  |  |
| 5 | 21,853,363 | HOM |  |  |  | CA | C | DEL |  |  |  |  |
| 5 | 23,199,299 | **HOM** |  |  |  | C | A | SNP | stop_gained | AT5G57260 |  |  |
| 5 | 26,711,631 | HOM |  |  |  | CCTATGGA | C | DEL |  |  |  |  |

_HOM: homozygous, HET: heterozygous, SNP: single nucleotide polymorphism, DEL: deletion, INS: insertion, ITX: intra-chromosomal translocation, CTX: inter-chromosomal translocation_

**Supplementary Table 3:** Whole genome sequence analysis results of C30-81-as6-77 mutant.

| Chromosome 1 | Position 1 | Genetic homogenity | Chromosome 2 | Position 2 | Genetic homogenity | Reference | Mutated base | Type of mutation | Effect type for gene | Gene ID affected by position 1 | Gene ID affected by position 2 | Mutation ID |
| --- | --- | --- | --- | --- | --- | --- | --- | --- | --- | --- | --- | --- |
| 1 | 8764 | HET |  |  |  | C | A | SNP |  |  |  |  |
| 1 | 3093370 | HET |  |  |  | C | T | SNP |  |  |  |  |
| 1 | 3190735 | HET |  |  |  | T | TA | INS | frameshift_variant | AT1G09820 |  |  |
| 1 | 3190736 | HET |  |  |  | G | C | SNP | missense_variant | AT1G09820 |  |  |
| 1 | 7925164 | HET |  |  |  | TA | T | DEL |  |  |  |  |
| 1 | 10085618 | HET |  |  |  | G | A | SNP |  |  |  |  |
| 1 | 11639575 | HET |  |  |  | AT | A | DEL |  |  |  |  |
| 1 | 17740018 | HET |  |  |  | T | C | SNP |  |  |  |  |
| 1 | 18341659 | HET |  |  |  | T | G | SNP |  |  |  | (1) |
| 1 | 18557695 | HET |  |  |  | A | G | SNP |  |  |  | (2) |
| 1 | 20690572 | HET |  |  |  | T | A | SNP |  |  |  |  |
| 1 | 22,888,469 | **HET** | 2 | 10,120,869 | **HET** |  |  | CTX | trancation |  | AT2G23770 |  |
| 1 | 22,888,477 | **HET** | 2 | 10,120,491 | **HET** |  |  | CTX | trancation |  | AT2G23770 |  |
| 2 | 8682241 | HET |  |  |  | G | A | SNP |  |  |  |  |
| 2 | 10,630,044 | **HET** | 2 | 10,630,167 | **HET** |  |  | ITX |  |  |  |  |
| 2 | 10,630,186 | HET |  |  |  |  | 27 bp DEL | DEL |  |  |  | (3) |
| 2 | 11263038 | HET |  |  |  | T | A | SNP |  |  |  |  |
| 2 | 14123848 | HET |  |  |  | T | G | SNP |  |  |  |  |
| 2 | 16930529 | HET |  |  |  | AC | A | DEL |  |  |  |  |
| 2 | 19205018 | HET |  |  |  | AT | A | DEL |  |  |  |  |
| 3 | 1217043 | HOM |  |  |  | G | C | SNP |  |  |  |  |
| 3 | 1709369 | HOM |  |  |  | C | T | SNP | stop_gained | AT3G05760 |  |  |
| 3 | 5528102 | HOM |  |  |  | C | T | SNP | missense_variant | AT3G16310 |  |  |
| 3 | 7581255 | HOM |  |  |  | C | T | SNP |  |  |  |  |
| 3 | 11719665 | HOM |  |  |  | C | A | SNP |  |  |  |  |
| 3 | 13723518 | HOM |  |  |  | AGTTTGGCCTCATGGTCTC | A | DEL |  |  |  |  |
| 3 | 13723540 | HOM |  |  |  | G | A | SNP |  |  |  |  |
| 3 | 13723541 | HOM |  |  |  | T | A | SNP |  |  |  |  |
| 3 | 13723542 | HOM |  |  |  | T | A | SNP |  |  |  |  |
| 3 | 14315102 | HOM |  |  |  | C | T | SNP |  |  |  |  |
| 3 | 15186549 | HOM |  |  |  | GGTCAAAATTTTTTATTGCCATACCAT | G | DEL |  |  |  |  |
| 3 | 15186550 | HOM |  |  |  |  | 27 bp DEL | DEL |  |  |  |  |
| 3 | 16235901 | HOM |  |  |  | TTC | T | DEL |  |  |  |  |
| 3 | 18217794 | HOM |  |  |  | A | T | SNP | missense_variant | AT3G49142 |  |  |
| 3 | 19710233 | HOM |  |  |  | G | C | SNP |  |  |  |  |
| 3 | 22702225 | HOM |  |  |  | A | C | SNP | missense_variant | AT3G61340 |  |  |
| 4 | 347586 | HOM |  |  |  | T | C | SNP |  |  |  |  |
| 4 | 2913480 | HOM |  |  |  |  | 30 bp DEL | DEL |  |  |  |  |
| 4 | 8928591 | HET |  |  |  | G | A | SNP |  |  |  |  |
| 4 | 9298550 | HET |  |  |  | G | C | SNP |  |  |  |  |
| 4 | 10709765 | HET |  |  |  | C | A | SNP |  |  |  |  |
| 4 | 16359953 | HET |  |  |  | GA | G | DEL |  |  |  |  |
| 4 | 17509276 | HET |  |  |  | G | T | SNP | missense_variant | AT4G37190 |  |  |
| 5 | 8489292 | HOM |  |  |  | T | C | SNP | missense_variant | AT5G24740 |  |  |
| 5 | 23199299 | HET |  |  |  | C | A | SNP | stop_gained | AT5G57260 |  |  |
| 5 | 26711631 | HET |  |  |  | CCTATGGA | C | DEL |  |  |  |  |

_HOM: homozygous, HET: heterozygous, SNP: single nucleotide polymorphism, DEL: deletion, INS: insertion, ITX: intra-chromosomal translocation, CTX: inter-chromosomal translocation_
